# Plasma Cell-Free DNA in Acute and Chronic Aortic Syndromes

**DOI:** 10.1101/2025.01.06.631373

**Authors:** Ruwan A Weerakkody, Martyna Adamowicz, Christelle Robert, John Pound, Maaz Syed, Caroline Pumpe, Alsadeg Bilal, Alexander Laird, Duncan B McLaren, Orwa Falah, Andrew L Tambyraja, Matthew J Reed, David E Newby, Timothy J Aitman

**Affiliations:** Institute of Genetics and Cancer, University of Edinburgh, Edinburgh, UK; Department of Vascular Surgery, Royal Infirmary Edinburgh, Edinburgh, UK; BHF Centre of Research Excellence, University of Edinburgh, Edinburgh, UK; Institute for Regeneration and Repair, University of Edinburgh, Edinburgh, UK; Department of Urology, NHS Lothian, Edinburgh, UK; Edinburgh Cancer Centre, University of Edinburgh, Edinburgh, UK; Emergency Medicine Research Group Edinburgh (EMERGE), Royal Infirmary of Edinburgh, Edinburgh, UK; Acute Care Edinburgh, Usher Institute, University of Edinburgh, Edinburgh, UK

## Abstract

Acute aortic syndromes and other acute aortic events are commonly misdiagnosed and associated with high mortality and morbidity. There is need for a blood-based biomarker to indicate the acutely unstable aorta. Plasma cell-free DNA (cfDNA) is an established diagnostic marker in prenatal testing and cancer. Its use as a potential diagnostic marker in aortic disease remains to be demonstrated.

We found that plasma short-fragment (50-700bp) cfDNA concentration is significantly increased in patients with both acute and chronic aortic disease compared with several control groups, and is significantly increased in acute versus chronic aortic disease. Plasma sf-cfDNA may differentiate acute aortic syndromes from other acute presentations of chest pain with good diagnostic accuracy and could prove a significant advance in the management of aortic disease.

## Introduction

Acute aortic syndromes (AAS) - acute aortic dissection, intra-mural haematoma and penetrating aortic ulcer - are increasing in incidence. AAS, and other acute aortic events including ruptured aneurysm and aneurysms at imminent risk of rupture, are frequently misdiagnosed and associated with high morbidity and mortality.^1^ Timely diagnosis and treatment improves clinical outcomes.^1^ Computed tomographic angiography (CTA), currently the gold standard diagnostic test, is not practicable as a routine screening method in the emergency department (ED) due to resource implications, low diagnostic yields and exposure to ionising radiation. Further, CTA is in itself unable to reliably predict imminent aortic complications. Blood-based biomarkers to aid diagnosis and to enable prediction of high-risk aortic events represent an urgently needed point-of-care diagnostic tool.^1,2^

Analysis of circulating cell-free DNA (cfDNA), also known as “liquid biopsy”, is a clinical diagnostic methodology that has transformed prenatal genetic testing, is entering routine practice in cancer management and has more recently shown promise as a biomarker of tissue damage in cardiovascular disorders.^3,4^ Its use as a potential diagnostic marker in aortic disease remains to be demonstrated.

## Methods

Here, we quantified plasma cfDNA in two prospectively recruited cohorts of patients with aortic disease: an “Acute Aortic” cohort comprising AAS, symptomatic aortic aneurysms and ruptured aortic aneurysms sampled within two weeks of symptom onset and before surgical intervention; and a “Chronic Aortic” cohort comprising AAS in the chronic asymptomatic phase of disease and chronic asymptomatic thoracic, abdominal and thoraco-abdominal aortic aneurysms all sampled at routine outpatient review more than 3 months from initial presentation. These were compared with prospective control cohorts with a range of conditions including: patients presenting to ED with acute non-aortic chest pain, outpatients with uncomplicated varicose veins, healthy volunteers and patients with pre-treatment kidney or prostate cancer. Apart from the two cancer cohorts, patients with active malignancy or inflammatory states were excluded. Informed consent followed approved protocols from the East of Scotland and Edinburgh Medical School Research Ethics Committees (Ref 20/ES/0061 projects SR1968, SR1514, SR1040; and EMREC Ref 21-EMREC-041).

Blood samples from all cohorts and control groups was collected into PAXgene® Blood cfDNA tubes, enabling room temperature storage for several days without leukocyte or red cell lysis. Plasma was separated by centrifugation within 4 days and plasma cfDNA extracted using a modification of the QIAmp Circulating nucleic acid/QIAmp MinElute virus spin protocols (Qiagen) to optimise DNA yield. cfDNA was quantified by Qubit (Invitrogen) and Qubit dsDNA HS Assay (ThermoFisher) and the percentage of short-fragment cfDNA (sf-cfDNA; 50-700 bp) quantified by TapeStation 4200 (Agilent). As validation of the Qubit/Tapestation methodology, sf-cfDNA was also measured by the orthogonal method LINE-PCR,^5^ with strong correlation between the two methods for measurement of sf-cfDNA (r^2^=0.87; p<0.05). Significance differences in plasma sf-cfDNA concentration between clinical groups were determined by two-tailed Mann-Whitney U test.

## Results

Plasma sf-cfDNA concentration was markedly higher in patients with acute aortic presentation than in the control populations and was also higher than in patients with chronic aortic disease (Figure 1A). To evaluate the performance of plasma sf-cfDNA concentration in discriminating between relevant pairs of sample types within this cohort data, Receiver Operating Characteristic (ROC) curves were constructed using the R package pROC, based on the ranking of sf-cfDNA concentrations in descending order and systematically applying thresholds to calculate sensitivity and specificity. The ROC analyses (Figure 1B and 1C) indicated that plasma sf-cfDNA may serve as a highly sensitive and specific marker for differentiating acute aortic syndromes from other acute causes of chest pain (area under the curve, mean AUC = 0.90, Figure 1B) and as a moderate discriminator of acute versus chronic aortic disease (mean AUC = 0.66, Figure 1C). However, as this evaluation is based solely on this single dataset, the performances should be interpreted cautiously with future validation in larger cohorts required to confirm generalisability.

**Figure 1.**
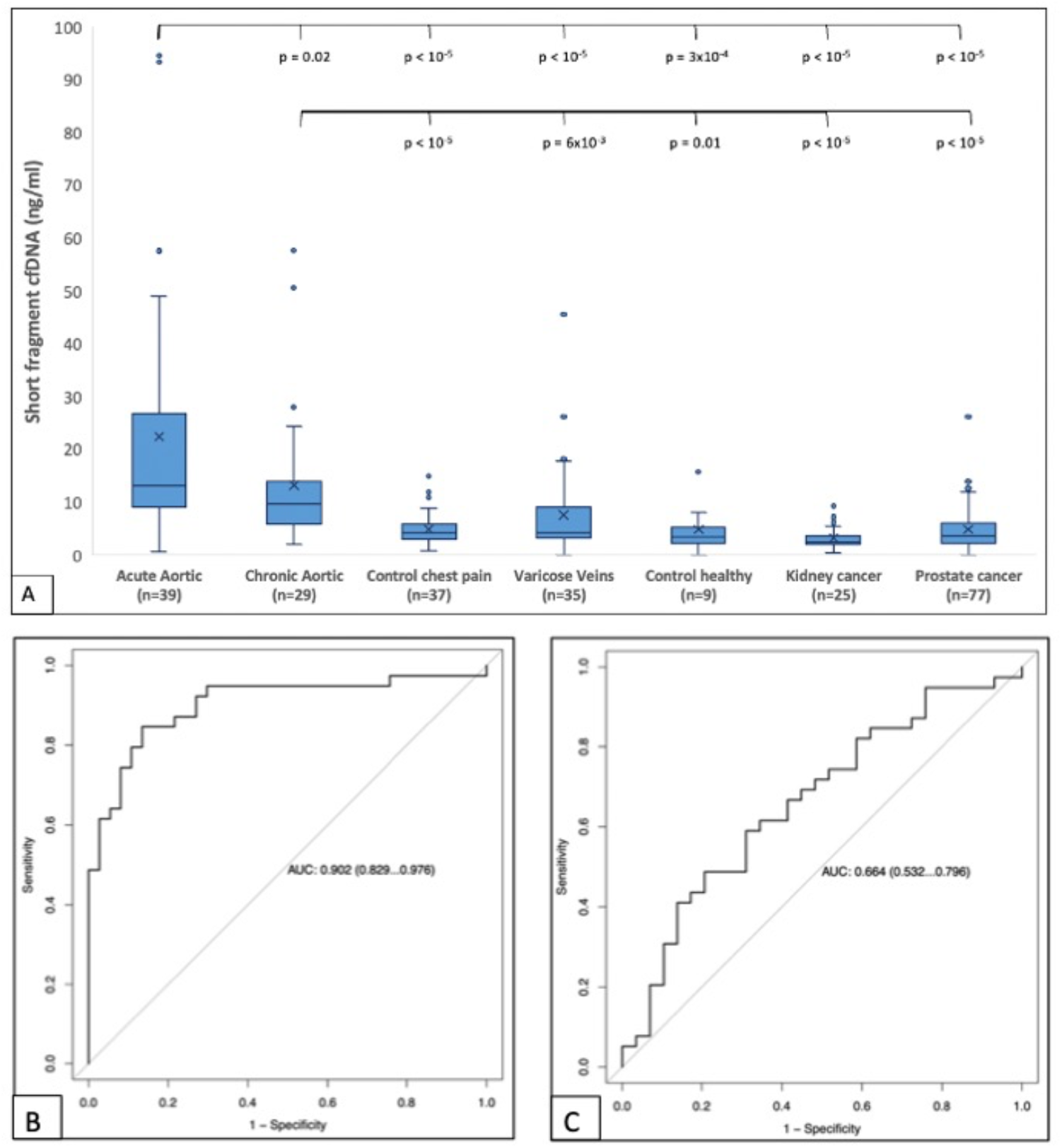
**Panel A: Box-plots** of Plasma short-fragment cfDNA yield (ng/uL) by clinical subgroup: 1. Acute Aortic cohort (as defined in text), 2. Chronic Aortic cohort (as defined in text), 3. Control-chest pain (prospective cohort of acute non-aortic chest pain presenting to ED), 4. Varicose veins (prospective cohort of uncomplicated varicose vein outpatients), 5. kidney cancer (newly diagnosed kidney cancer patients prior to commencement of treatment, various histological types), and 6. prostate cancer (newly diagnosed prostate cancer patients, prior to treatment or surgery). Significant differences between clinical groups were determined by two-tailed Mann-Whitney test (p-values indicate comparisons with the Acute Aortic cohort (upper row of p-values) and with the Chronic Aortic cohort (lower row of p-values)). **Panels B & C:** Receiver Operating Characteristic curves estimating the sensitivity and specificity of short-fragment cfDNA assays. (**B)** Assay performance in distinguishing Acute Aortic patients (N=39) from Control-Chest Pain cohort (N=37). The mean AUC of 0.902 indicates a model with strong discriminative performance. **(C)** Assay performance in distinguishing Acute Aortic patients (N=39) from Chronic Aortic patients (N=29). The mean AUC of 0.664 indicates a model with performance above a random classifier, though with wide 95% confidence intervals (0.532-0.796). 95% confidence intervals were computed with 2000 stratified bootstrap replicates.

In this cohort, we found no significant correlation between maximum aortic diameter and plasma sf-cfDNA concentration in either aortic cohort (Acute Aortic: mean maximum diameter 48mm, range 26-90mm; Chronic Aortic: mean maximum diameter 59mm, range 30-100mm).

## Conclusion

Our findings suggest that plasma sf-cfDNA may differentiate acute aortic syndromes from other acute presentations of chest pain with good diagnostic accuracy, but that distinction of the acutely unstable aorta from the chronic stable state requires an assay with greater specificity. Further work is required to validate these data in larger cohorts and to refine assay specificity in patients with chronic aortic disease, for example by tissue-of-origin methylation analysis of plasma cfDNA. If thus replicated, cfDNA analysis could prove a significant advance in the management of aortic disease.

